# Evolutionary dynamics and ecological influences on Felidae

**DOI:** 10.1101/2023.06.29.547091

**Authors:** Lucas M. V. Porto, Gabriel P. de Olveira, Maico S. Fiedler

## Abstract

In the field of evolutionary biology, researchers have long been interested in comprehending the patterns of diversification and elucidating the mechanisms responsible for the impressive variety of lifeforms observed within taxonomic groups. Based on the fossil records of Felidae, and using a Bayesian framework, we assess here how speciation and extinction rates vary across the last 25 Myr, and how it can potentially be associated with changes in two major ecomorphological traits of Felidae, body mass and carnassial teeth, over such time span. We found two coupled independent increases in both traits along Felidae evolution, one in Machairodontinae, which gradually increases over the last 15 Myr, and the second in *Panthera*, which rapidly increases within 5 million years. Furthermore, gradual reductions in body mass and carnassial teeth were also observed for some genera such as *Leopardus, Lynx*, and *Felis*. Diversification rates showed a gradual reduction throughout the Miocene and Pliocene, with significant rate shifts occurring around 9 and 2 Myr. We suggest that felids were influenced by major environmental changes over the last 17 Myr that probably shaped both traits studied here in order to hunt larger herbivores, but also to explore new niches that became available.

## INTRODUCTION

Understanding the patterns of diversification is a central question in evolutionary biology (Rabosky 2010; Slater and Friscia 2019). Over the decades, biologists seek to uncover the mechanisms driving the remarkable diversity observed across different clades and identify the underlying attributes that contribute to the heterogeneous patterns of species richness (Slater 2015; Beaulieu and O’Meara 2016; Springer et al. 2019; Porto et al. 2023). The availability of diverse ecological niches can provide fertile ground for species to evolve unique traits and exploit new resources, leading to increased diversification rates (Figueirido et al. 2015; Porto et al. 2023). Additionally, selective pressures, such as competition for limited resources or interactions with other species, can drive the evolution of specialized traits and promote character displacement (Silvestro et al. 2015; Pires et al. 2017). However, biotic and abiotic factors not only can foster diversification but may also hinder it, leading to the decline of whole clades. By unraveling the complex interplay of these factors, scientists gain valuable insights into the drivers of species diversification observed in nature.

Phylogenetic studies are a key tool in understanding the evolutionary relationships among traits and diversification rates. By analyzing molecular data, researchers can reconstruct the Tree of Life and uncover how species are related to one another. However, relying solely on molecular phylogenies based on extant species can lead to incomplete or inaccurate reconstructions of evolutionary history (Quental and Marshall 2010). Without accounting for fossil data, researchers risk overestimating divergence times and having an incomplete understanding of evolutionary relationships (Quental and Marshall 2010; Wiens et al. 2010; Pyron 2015). As such, it is crucial to incorporate both molecular and fossil data into phylogenetic analyses to obtain a more accurate and comprehensive view of past events on clades’ evolution.

Felids represent a diverse group of carnivores comprising various species across the globe and have long fascinated researchers due to their remarkable morphological and ecological adaptations (Wilson and Mittermeier 2009). One intriguing aspect of felid evolution is a wide range of body masses that we can observe, not only on the fossil record, but in the extant species too, from large apex predators with 300kg to smaller, more specialized hunters ranging around 1kg (Sunquist and Sunquist 2002; Cuff et al. 2015).

Felids, akin to other carnivorous species, exhibit certain specializations that enhance their predatory behavior. One such adaptation is the presence of carnassial teeth, a distinctive shearing structure observed across all members of the Carnivora order (Werdelin et al. 2010). It is hypothesized that the evolution of specialized dental structures like the carnassial teeth, in conjunction with the diversity in body mass, played a significant role in felid diversification (Christiansen 2008; Meachen-Samuels and Van Valkenburgh 2009; Slater and Van Valkenburgh 2009; Pires et al. 2017). The forces behind such evolutionary adaptations are thought to be related to the availability of prey and inter-species competition, both of which are key drivers in the evolution of the Felidae family (Meachen-Samuels and Van Valkenburgh 2009; Pires et al. 2017).

By integrating a molecular phylogeny with a robust dataset of fossil species, and employing phylogenetic comparative methods, we aim to investigate the long-term trends in felid diversification and its potential association with body mass and carnassial teeth over the past 25 million years. We hypothesize that changes in the diversification of Felidae will be accompanied by variations in body mass and carnassial teeth. Studies have suggested that shifts in climate and vegetation over the past 30 million years have played a key role in driving diversification rates in mammals, with periods of environmental instability often coinciding with major changes in diversification rates (Werdelin and Peigné 2010; Badgley et al. 2017). Thus, we argue that, as Earth experienced major environmental changes over the last 25 Myr, such as the Middle Miocene Climatic Optimum, a geological event associated with climatic and paleogeographic changes that impacted mammals across the Northern Hemisphere (Zachos et al. 2008; Eronen et al. 2012; Casanovas-Vilar et al. 2014), novel environments and niches could have selective and evolutive pressures for the diversification of felids.

## MATERIAL AND METHODS

### Data collection

We used the robust Felidae phylogeny from Piras et al. (2013), which incorporates both extant and extinct species, presenting 35 of all the 37 extant species, and 59 extinct Felidae. Subsequently, we obtained the dataset of body mass and dental attributes from the *CarniFOSS* dataset (Faurby et al. 2021). We searched for all Felidae species in this dataset and compared them with the phylogeny from Piras et al. (2013). Any Felidae species present in the *CarniFOSS* dataset but not included in the phylogeny were added through phylogenetic imputation. Specifically, these species were placed at the root of their respective genera using the *Phytools* package (Revell 2012) in *R 4*.*3*.*0* (R Development Core Team 2023). On the other hand, Felidae species present in the phylogeny but not found in the *CarniFOSS* dataset were removed from further analysis, so evolutionary rates and trait reconstruction would be estimated with the same number of species.

With all selected species, we performed molecular dating to estimate the divergence times within the Felidae phylogeny. We incorporated the known node ages from Piras et al. (2013) as well as information from the *TimeTree* database (Kumar et al. 2022) for those species added to the phylogeny. This resulted in a non-ultrametric phylogeny consisting of 71 species (14 extant and 57 extinct), with 30 felids that were imputed in the phylogeny (Figure 1 and Tables S1). Time-calibration was performed using the *chronopl* function from the *Ape* package (Paradis and Schliep 2019), which is a penalized likelihood method to estimate node ages following Sanderson (2002). For the penalization process, we used λ (lambda) = 1.

**Figure 1.**
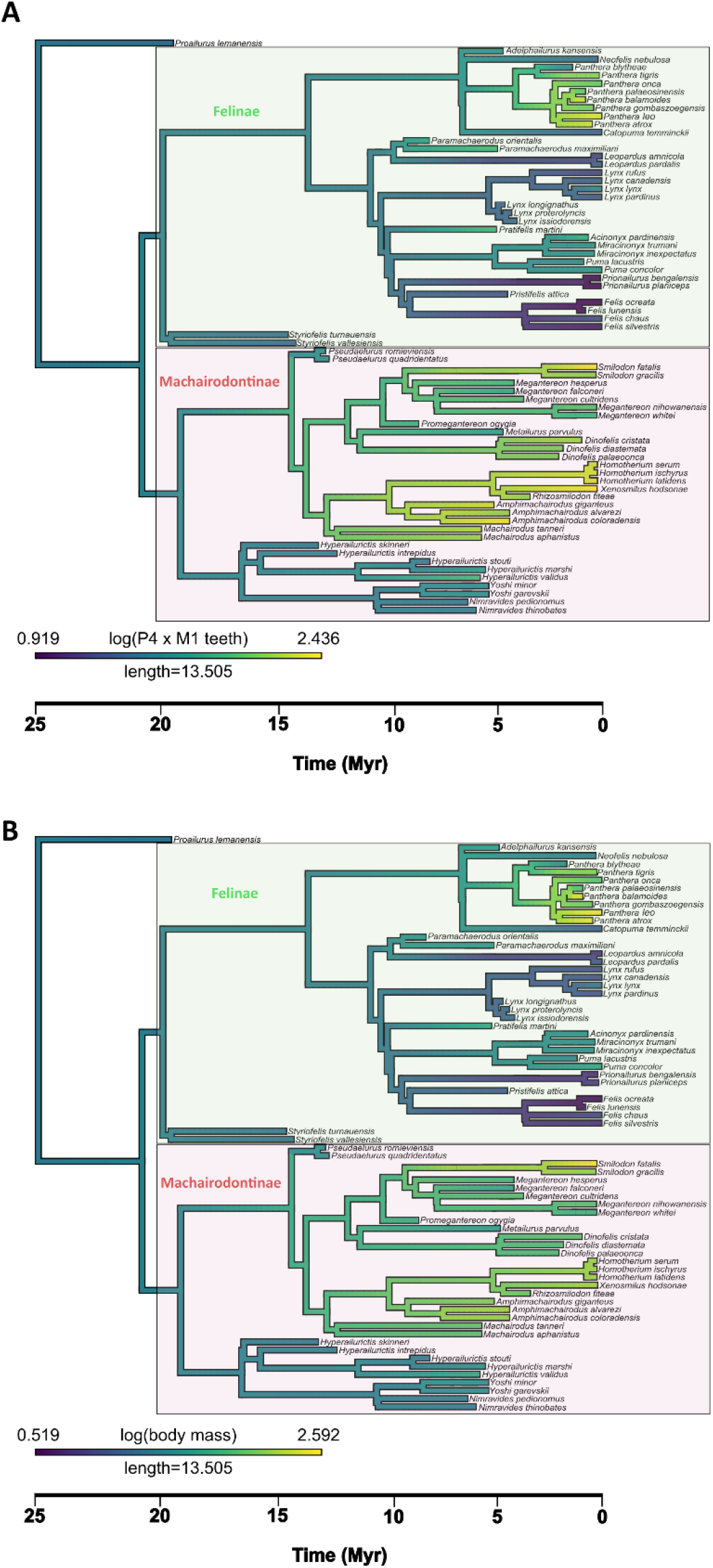
The phylogenetic tree used here with the reconstructed values of carnassial teeth (A) and body mass (B) over the evolution of Felidae. Such reconstructed trees were used to obtain time slices and get mean trait values over time. Green and red boxes indicate the Felinae and Machairodontinae subfamilies, respectively.

The phylogeny of Felidae is represented by two subfamilies, Machairodontinae and Felinae. The Machairodontinae clade is extinct and includes genera like *Smilodon* and *Homotherium* (Figure 1). Felinae, on the other hand, is an extant clade that includes all the living felids on the planet at the moment, being represented by genera such as *Panthera, Felis*, and *Lynx* (Figure 1) (Piras et al. 2018).

We then used the *Paleobiology Database* (PBDB 2023) to retrieve fossil occurrence records for these 71 species across different geographic locations worldwide (Figure S1). The data collection process from *PBDB* yielded a comprehensive dataset comprising 588 fossil records (Appendix 1). The data were downloaded on June 3rd, 2023, using the group name “Felidae” and the following parameters: Type = Occurrences, Taxonomic resolution = species (accepted names only), time intervals = all, region = all continents.

From *CarniFOSS* we used the logarithmic scale data of body mass (g) and the length (mm) of the upper fourth premolar (P4) and lower first molar (M1) (Table S1). P4 and M1 are responsible for the carnassial teeth in Felidae (Wilson and Mittermeier 2009). In order to have a measure that would be a proxy for the shape of the carnassial teeth and a good indicative of the eco-morphology of felines, the use of the Relative Blade Length (trigonid blade length relative to the total length of the lower first molar), also known as RBL, would be the best approach (Van Valkenburgh 1989). However, to get RBL values, one needs a complete description of the structures of P4 and M1, which are not available for all the species we are using here. As such, we utilized the product of P4 and M1 measurements as a proxy for assessing the size of the carnassial teeth in Felidae. This method draws on the strong correlation between RBL values and M1 tooth length in Carnivora as identified by de Bonis et al. (2009).

### Phylogenetic analysis

Our phylogenetic analyses consisted of two parts. Firstly, we performed an Ancestral Character Reconstruction for body mass and for our proxy for the size of the carnassial apparatus (P4 x M1). To accomplish this, we employed the *contmap* function from the *phytools* package (Revell 2012). This approach utilized maximum likelihood methods to infer the most likely ancestral states based on the observed character data and the branch lengths of the phylogeny.

The second part of our analysis focused on slicing over time both reconstructed phylogenies into intervals of 0.5 million years and calculating the mean value of each trait within those intervals. This step was crucial for capturing the temporal dynamics of body mass and the carnassial apparatus, considering that our phylogeny was not ultrametric. By partitioning the phylogenies into uniform time intervals, we could assess how the average body mass and carnassial teeth varied over time based on the lineages present within each interval. The *R* code developed for this part is available in the supplementary material.

### Diversification rates

We estimated species speciation and extinction rates, and consequently, the net diversification rate, using fossil records from *PBDB* for all the species in the phylogeny constructed here adapted from Piras et al. (2013). We conducted our analysis using the *PyRate* software (Silvestro et al. 2014), a Bayesian framework that integrates preservation and diversification processes while accounting for the incompleteness of the fossil record. *PyRate* utilizes the Reversible Jump Birth-Death Markov Chain Monte Carlo algorithm to infer speciation and extinction rates and detect significant rate shifts through time (Silvestro et al. 2014).

Our analyses employed the *mG* argument, which is a Gamma model of rate heterogeneity, which enables us to account for heterogeneity in the preservation rate across lineages. We also used the *qShift* model that allows for the preservation rates to be estimated independently within each geological epoch. We set 30.000.000 iterations, sampling every 10.000 iterations to get posterior parameter estimates. We discarded the first 10% as burn-in. The time windows used to estimate the preservation rates were based on the distribution of fossil records over time (Figure S2). To optimize the concentration of fossil records in distinct intervals, our time boundaries were set at 28.5, 13.5, 5.3, 2.58 and, 0 million years ago.

To address the uncertainty in fossil occurrences, we generated 25 temporal replicates by randomly assigning ages within each occurrence timespan. *PyRate* analyses were conducted on each replicated dataset. To avoid pseudo-replicates, we used the “collection number” identifier from *PBDB* to identify occurrences from the same assemblages.

## RESULTS

### Phylogeny and ancestral states

The ancestral character reconstruction of body mass and carnassial teeth (Figures 1A and 1B) revealed strongly similar patterns in the evolutionary history of Felidae. In the case of body mass, the reconstruction indicated that the emergence of large sizes occurred independently in two distinct clades. Firstly, the Machairodontinae subfamily exhibited a gradual increase in body mass over the past 15 million years. On the other hand, within the Felinae subfamily, the genus *Panthera* displayed a remarkably rapid increase in body mass within a time span of just 5 million years.

Regarding the evolution of the carnassial teeth, the reconstruction indicated a similar trend. Large apparatus sizes appeared independently twice in the phylogeny, aligning with the emergence of big felids. The subfamily Machairodontinae exhibited a gradual and continuous increase in apparatus size over the same 15-million-year period, while *Panthera* experienced more rapid and concentrated development of large apparatus. The ancestral reconstruction also revealed two independent events of gradual reduction of body mass and carnassial teeth in two extant genera, *Felis* and *Leopardus*, over the past 10 million years.

Our analysis of the average values for body mass and carnassial size across the reconstructed tree revealed intriguing patterns in the evolution of Felidae (Figures 2B and 2C). Both attributes displayed dynamic changes over time. Between 17.5 and 9 million years ago, there was a notable increase in both body mass and carnassial teeth, suggesting a trend towards larger sizes within Felidae. However, this upward trajectory is interrupted between 9 and 2 million years ago, as both traits experience a pronounced decline until recent times.

**Figure 2.**
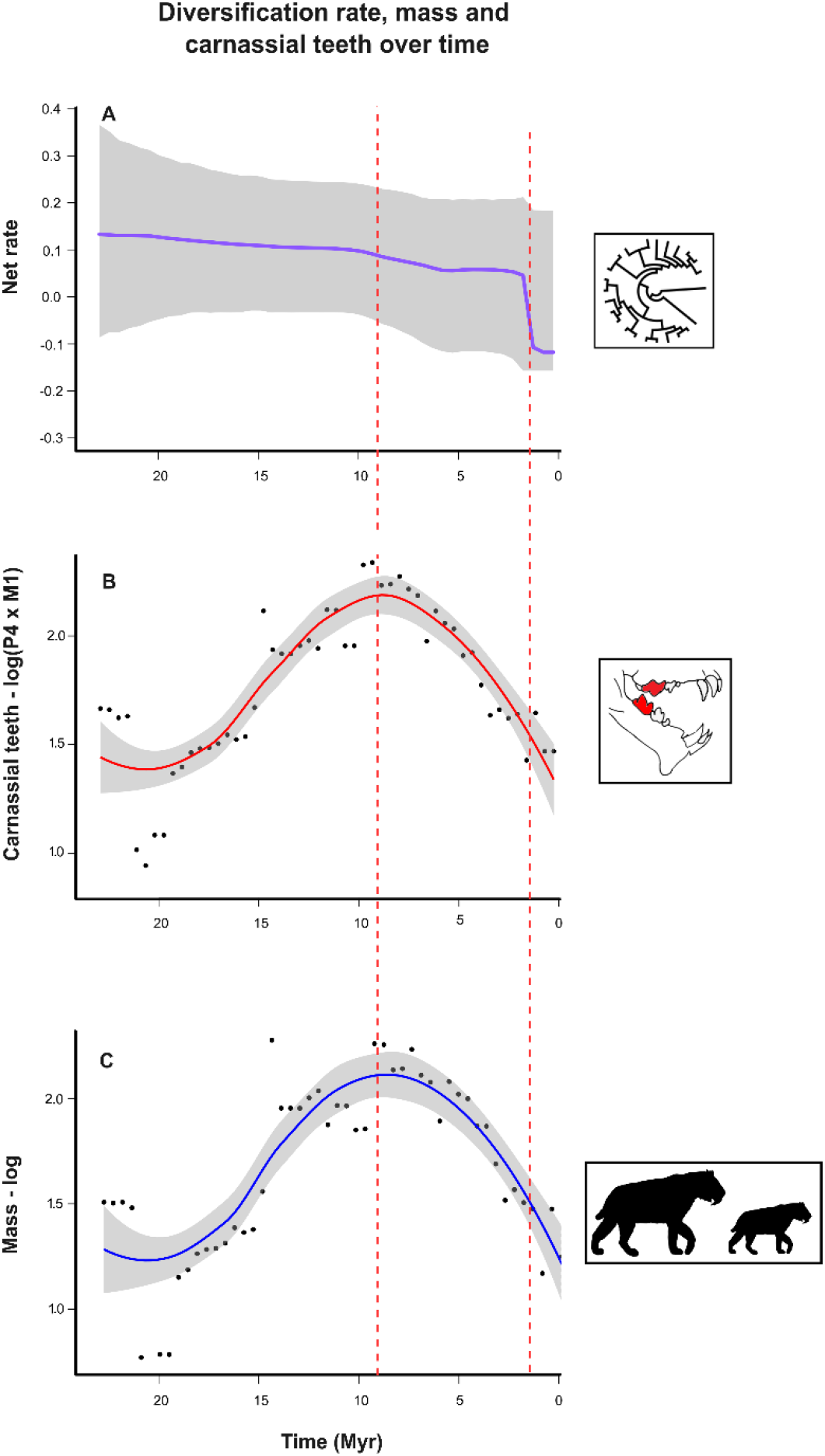
Panel showing the variation over time of the net diversification rate (A), reconstructed values of carnassial teeth (B), and reconstructed values of body mass (C). Black dots in plots B and C are the mean trait values of the 0.5 Myr time slices. Red dashed lines represent the moments in time of the two significant rate shifts in the diversification rate.

### Felidae diversification

*PyRate* analyses indicated two significant rate shifts during the diversification of Felidae, one around 9 million years ago and another around 2 million years ago (Figure 2A). Looking at the diversification rate (Figure 2A), our results showed a gradual reduction in the diversification of Felidae throughout the Miocene and much of the Pliocene, becoming a negative rate only around 2 million years ago, during the Late Pliocene. When we examine the rates of speciation and extinction separately, our results demonstrated that both traits exhibited distinct dynamics over time (Figures 3A and 3B). The speciation rate remained relatively constant over the past 25 million years, but the extinction rate showed an increase between 25 and 2 million years ago, followed by a sharp rise in a short period of time thereafter.

**Figure 3.**
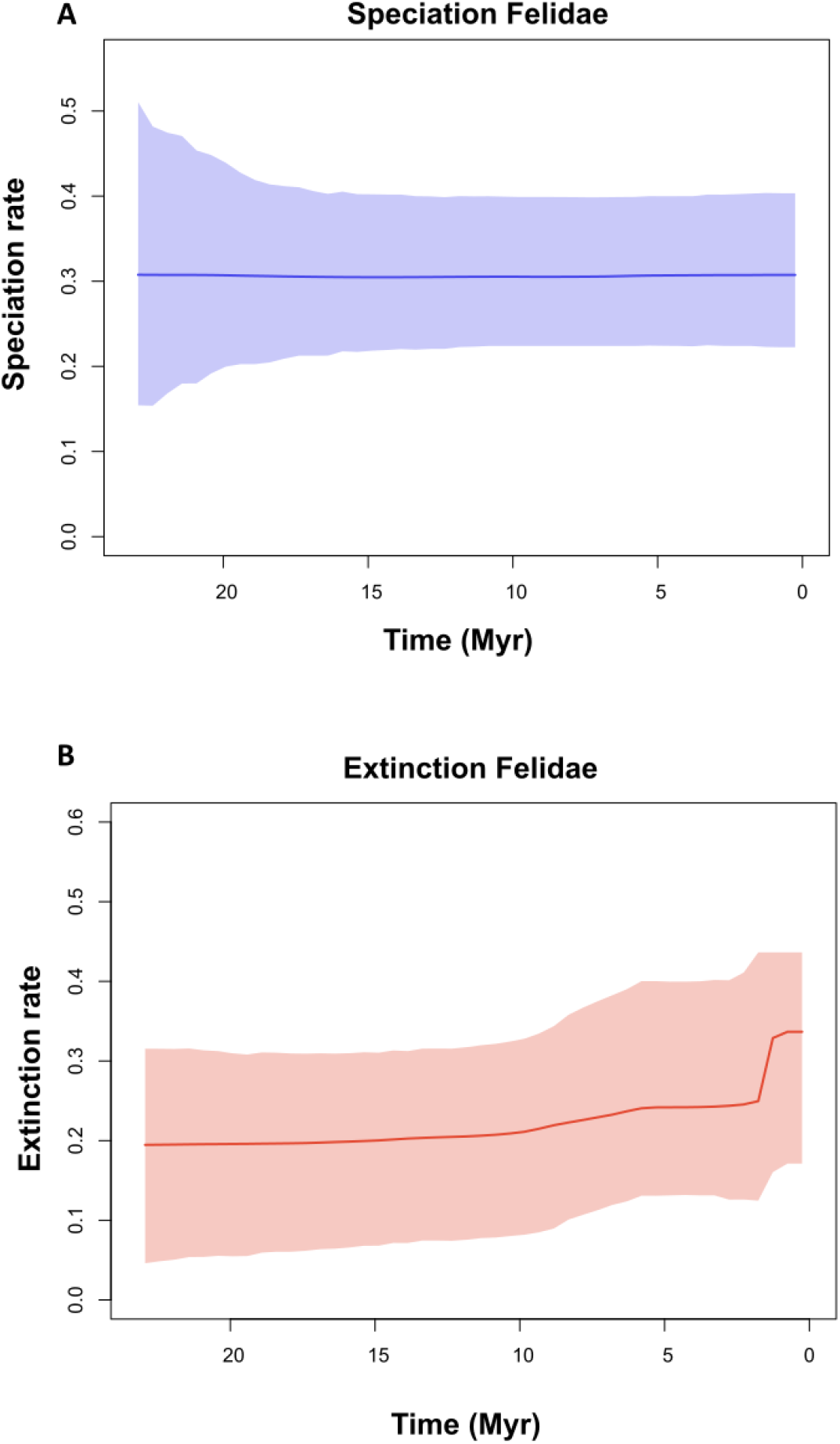
Speciation (A) and extinction (B) rates of Felidae evolution over the last 25 million years.

It is important to highlight that uncertainty remained stable over the whole estimates of diversification, extinction, and during most part of speciation, including the moments of significant rate shifts detected by *PyRate*.

## DISCUSSION

Although there was no linear correlation between ecomorphological traits and net diversification (Figures S3A and S3B), our results identified a moment in time when a significant decrease in the diversification rate of Felidae was followed by a major decline in medium values of body mass and carnassial teeth over time (Figure 2). In addition, our study revealed parallel trends in the evolution of body mass and carnassial teeth, presenting strongly similar variations over time, but also revealing independent events of increase and decrease trait values in distinct clades.

We observed a major increase in body size and carnassial teeth of Felidae lineages from 17 to 10 Myr, as indicated by the mean trait values measured across reconstructed phylogenies (Figures 2B and 2C). This pattern was particularly evident when examining the reconstructed phylogenies, with the Machairodontinae clade playing a significant role in driving this trend. The Machairodontinae clade developed stronger bite and bigger dental structures (Van Valkenburgh 2007; Randau et al. 2013). The rise of such specialized dental morphology likely evolved in Felidae because it provided advantages in competing for prey, particularly megaherbivores (Cantalapiedra et al. 2011; Van Valkenburgh et al. 2015). Although this specialized behavior did not manifest in a marked increase in diversification rate, it did contribute to the evolution of larger felids with proportionally larger teeth.

The increase in extinction rate around 9 Myr, which is identified here as a significant rate shift in the diversification rate, matches in time with the posterior period of the Middle Miocene Climatic Optimum (MMCO), a geological event associated with climatic and paleogeographic changes in association with declining temperatures and increasing aridity that resulted in a massive mammal turnover in the Northern Hemisphere (Zachos et al. 2008; Eronen et al. 2012; Casanovas-Vilar et al. 2014). Such geological event has emerged as a plausible explanation for both the observed declines in felid diversification rates, as well as reductions in body size and carnassial teeth. In addition, our estimates of both speciation and extinction rates are in line with the rates proposed by Pires et al. (2015), which supports the pattern found here for the net diversification of Felidae.

When examining the reconstructed ancestral characters, we observed the rise of a new clade, represented by genera such as *Lynx* and *Felis* (called from now on as the small-cats clade), during this period after the MCCO, which subsequently underwent rapid diversification. It is within this clade that we noticed a gradual evolution of small body sizes and small carnassial teeth for the first time in the Felidae lifespan (Figures 2B and 2C). In addition, at the same time, we observed the clade that presents genera like *Smilodon, Dinofelis*, and *Machairodus* (called from now on as the big-cats clade) experiencing for a long time no diversification at all. However, the post-MCCO alone may not explain it all, and an interplay between such geological event associated with the rise of pounce-pursuit canids in Europe (Wang and Tedford 2008; Figueirido et al. 2015), were probably responsible for the observed phylogenetic and morphological patterns described here.

There is some evidence to support the idea that the small-cats clade evolved small sizes to avoid competition with medium and large canids over the last 10 million years. Carnivore species tend to partition their prey resources based on body size, with larger predators targeting larger prey, and smaller predators targeting smaller prey (Carbone et al. 2007). This suggests that competition between felids and canids may have played a role in driving the evolution of both body mass and carnassial teeth in the small-cats clade, as smaller sizes would allow them to exploit different niche spaces and avoid direct competition with larger canid species. Such a trend could have allowed felids to hunt more efficiently in open habitats by focusing on smaller prey such as rodents, with no need for big carnassial teeth (Meachen-Samuels and Van Valkenburgh 2009).

Over the last 12 Myr, the Machairodontinae clade, in its majority, was still presenting a trend to higher values of body mass and carnassial teeth, evolving several sabertoothed forms across the planet. It is intriguing that, even after developing bigger bodies and carnassial teeth, aspects that confer ecological advantages over competitors (Slater and Van Valkenburgh 2008; Cantalapiedra et al. 2011), the clade was completely extinct. A probable explanation is that specialization into such unique adaptations acquired by these felids, which were advantageous in a previous moment between 17-10 Myr, led them to an evolutionary dead-end (Mondanaro et al. 2017). Piras et al. (2018) showed that sabertoothed lineages, when we look at the fossil record, exhibit a higher extinction rate compared to others. The extensive morphological specializations required to become sabertoothed, coupled with their preference for large-sized prey, likely contributed to this high extinction rate. These findings are supported by the results of Cardillo et al. (2005), who demonstrated a positive relationship between large body size and the risk of extinction in mammals.

The two significant rate shifts found here in the diversification rate seem to have been caused by distinct factors. The abrupt decline in the extinct rate, which resulted in the pattern observed in the net rate of Felidae over the last 2.5 Myr (Figure 2A), was probably led by the significant climate fluctuations on Earth over the Pleistocene, which led to series of glacial periods known as the *Ice Age* (Batchelor et al. 2019). These climatic changes had a profound impact on ecosystems worldwide, being marked by extensive ice sheets and lower sea levels (Lisiecki and Raymo 2005; Batchelor et al. 2019). These shifts in climate and habitat availability could have had substantial consequences for felid populations, reducing the availability of large prey, mainly for the Machairodontinae clade, that were so well adapted to a narrow niche that might have had a limited ability to evolve into generalists forms (Day et al. 2016). Although the arrival of humans in Eurasia and the Americas over the past 10,000 years negatively impacted the last lineages of Machairodontinae (DeSantis et al. 2012), our results suggest that this impact was not greater than the previous disturbances during the early Pleistocene, as the extinct rate is stable during this time.

## CONCLUSION

Overall, the findings presented here provide some insights into the association between the evolution of Felidae and two major ecomorphological traits. We suggest that felids were influenced by major environmental changes over the last 17 Myr, leading to a turnover of vegetation types and fauna across the Northern Hemisphere that probably shaped their body masses and dental adaptations in order to hunt larger herbivores that arise at this moment, but also to explore new niches that became available for small carnivores.

## TABLE LIST

**Table S1.**
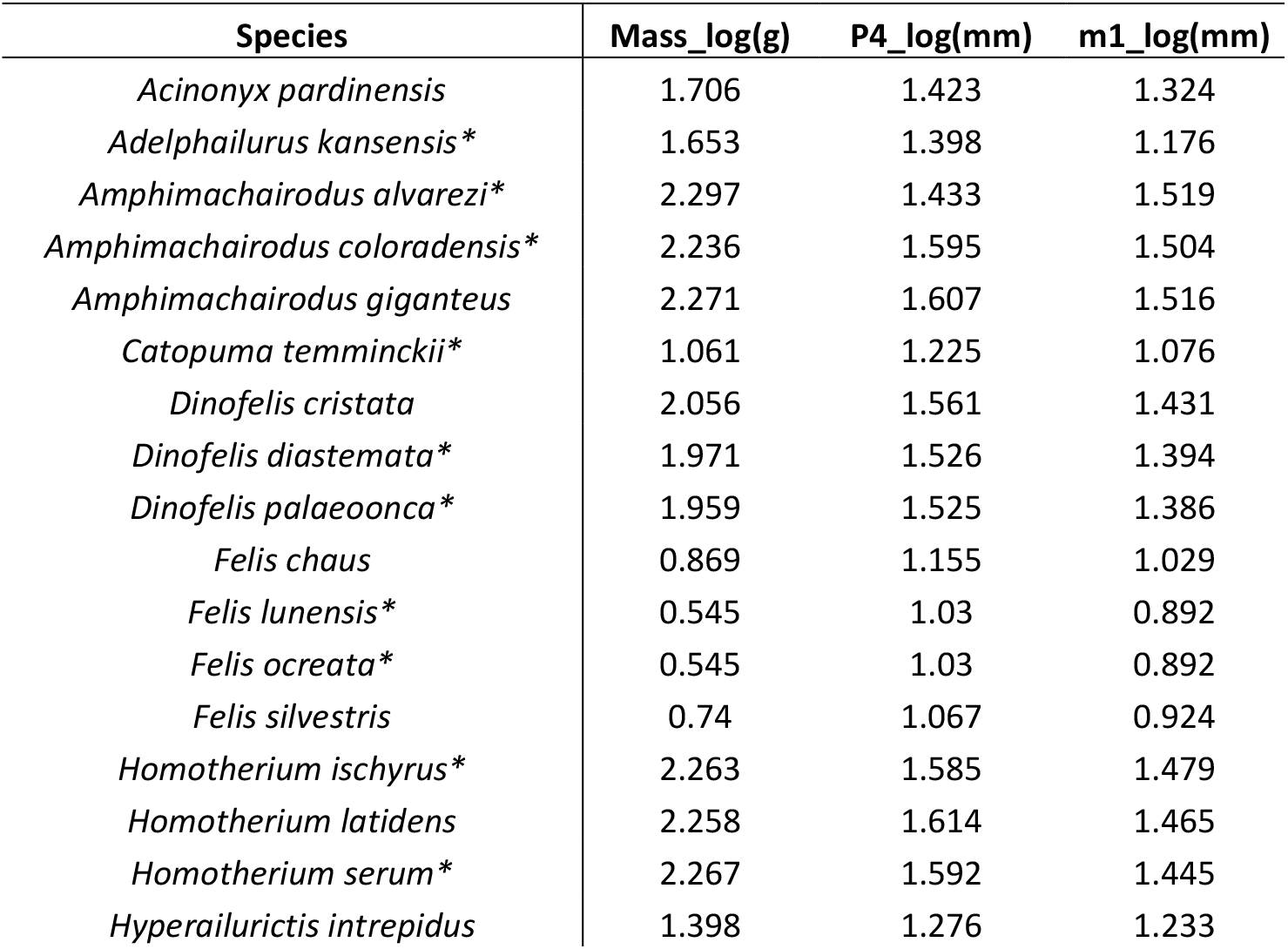
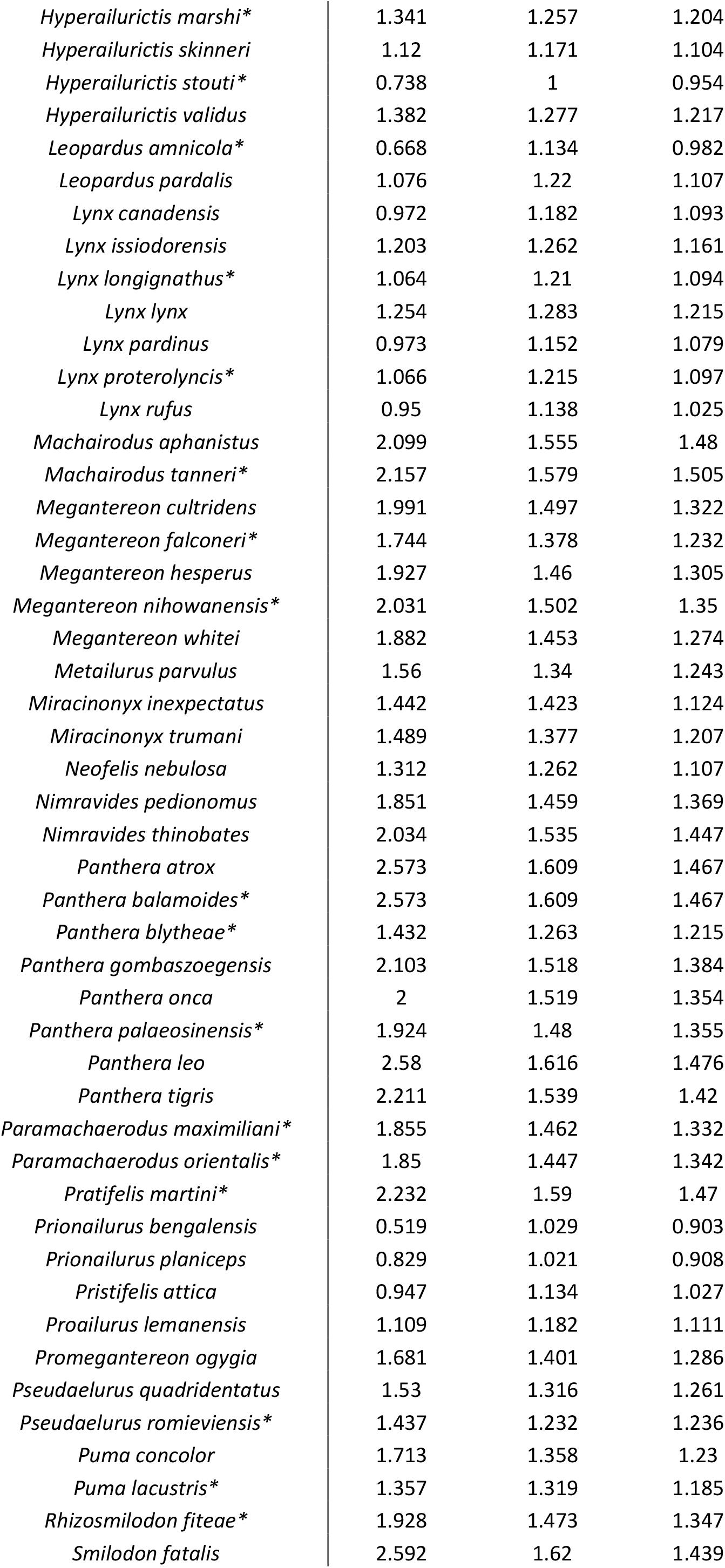
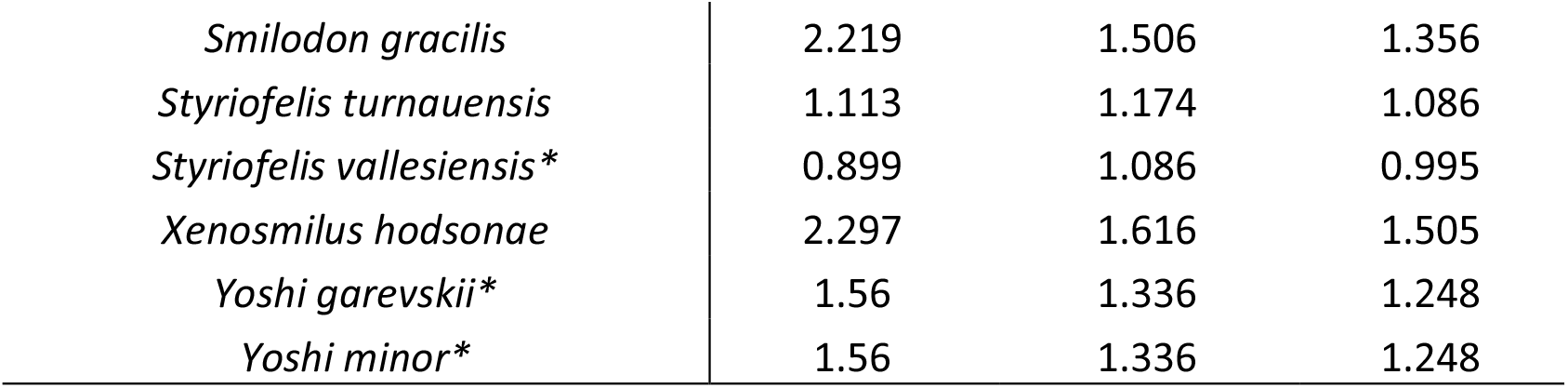
List of all the 71 Felidae species used here. Values of body mass and carnassial teeth (P4 and M1) are indicated. Species marked with ^*^ are the 30 felids that were imputed in the phylogeny from Piras et al. (2013).

## SUPPLEMENTARY MATERIAL

**Figure S1.**
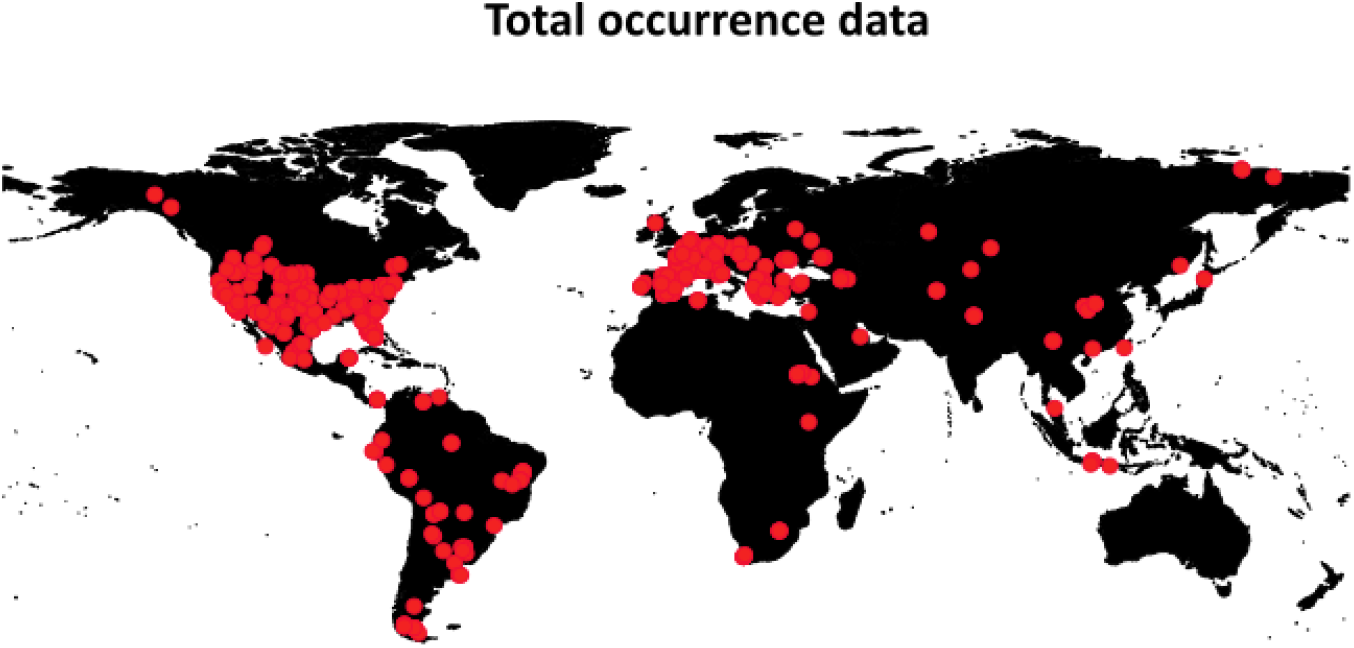
Map indicating the locations of the 588 fossil records from *PBDB* used here to estimate speciation and extinction rates.

**Figure S2.**
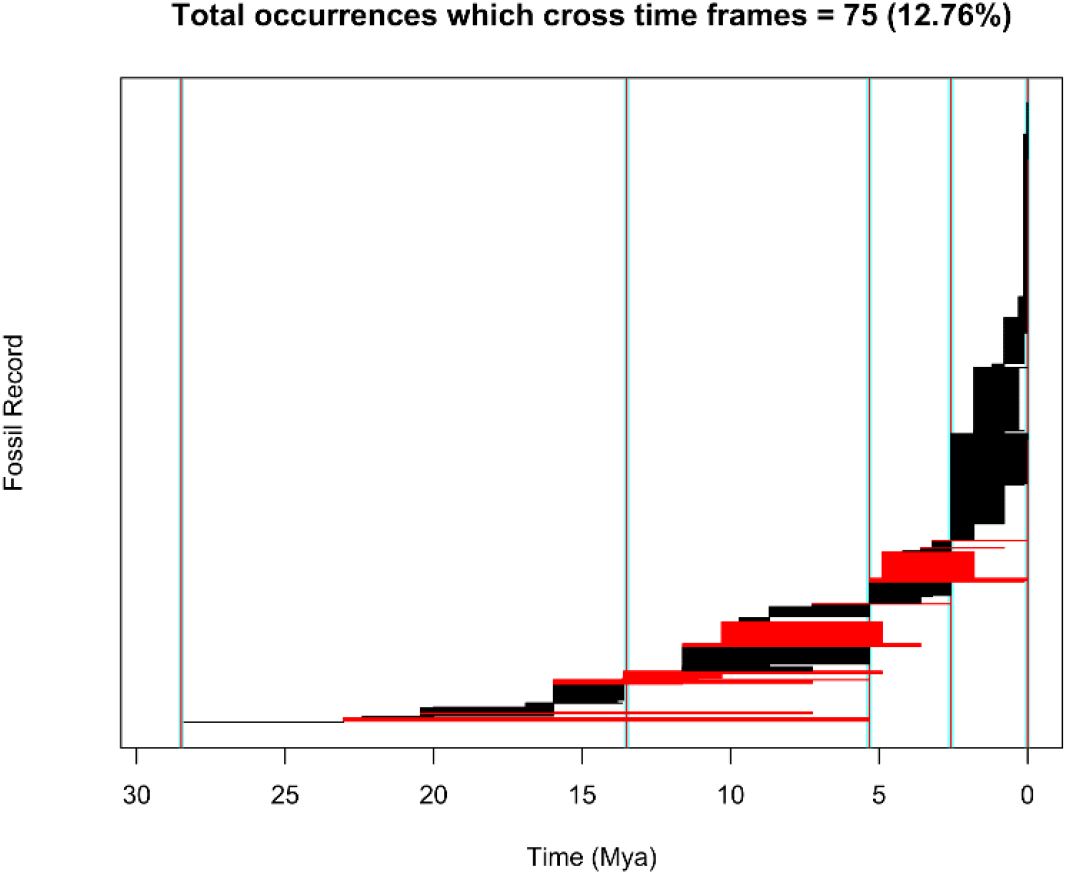
The time windows used to estimate the preservation rates used here (28.5, 13.5, 5.3, 2.58, and 0 Myr). Each line is a fossil record. Red lines are the occurrences that cross any of the time windows.

**Figure S3.**
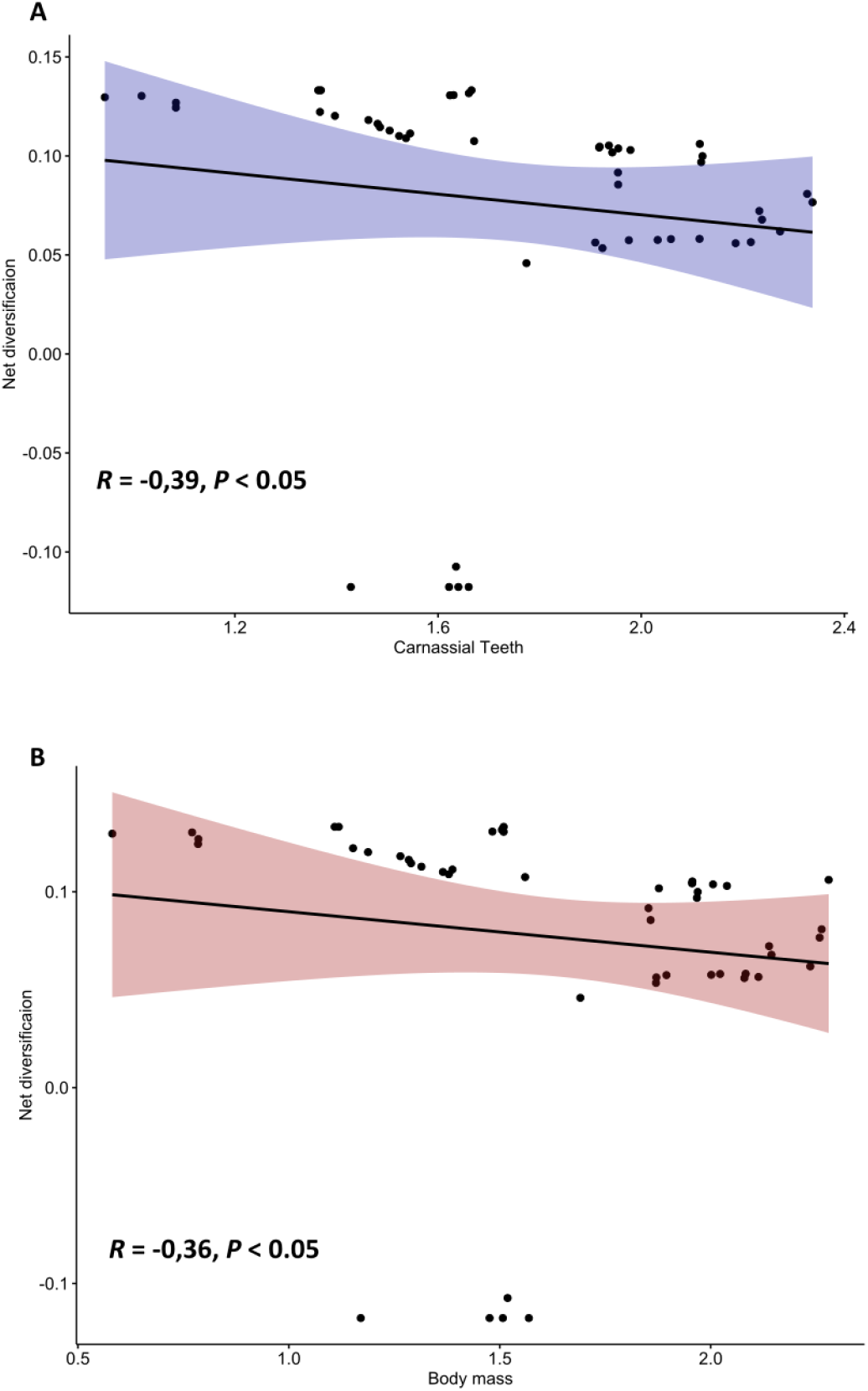
Correlation between the net diversification rate with the mean trait values of carnassial teeth (A) and body mass (B).

## REFERENCES

Badgley, C., T. M. Smiley, R. Terry, E. B. Davis, L. R. G. DeSantis, D. L. Fox, S. S. B. Hopkins, T. Jezkova, M. D. Matocq, N. Matzke, J. L. McGuire, A. Mulch, B. R. Riddle, V. L. Roth, J. X. Samuels, C. A. E. Strömberg, and B. J. Yanites. 2017. Biodiversity and Topographic Complexity: Modern and Geohistorical Perspectives. Trends Ecol. Evol. 32:211–226.

Batchelor, C. L., M. Margold, M. Krapp, D. K. Murton, A. S. Dalton, P. L. Gibbard, C. R. Stokes, J. B. Murton, and A. Manica. 2019. The configuration of Northern Hemisphere ice sheets through the Quaternary. Nat. Commun. 10:3713.

Beaulieu, J. M., and B. C. O’Meara. 2016. Detecting Hidden Diversification Shifts in Models of Trait-Dependent Speciation and Extinction. Syst. Biol. 65:583–601.

Cantalapiedra, J. L., M. Hernández Fernández, and J. Morales. 2011. Biomic Specialization and Speciation Rates in Ruminants (Cetartiodactyla, Mammalia): A Test of the Resource-Use Hypothesis at the Global Scale. PLoS One 6:e28749.

Carbone, C., A. Teacher, and J. M. Rowcliffe. 2007. The Costs of Carnivory. PLoS Biol. 5:e22.

Cardillo, M., G. M. Mace, K. E. Jones, J. Bielby, O. R. P. Bininda-Emonds, W. Sechrest, C. D. L. Orme, and A. Purvis. 2005. Multiple Causes of High Extinction Risk in Large Mammal Species. Science (80-.). 309:1239–1241.

Casanovas-Vilar, I., L. W. Van den Hoek Ostende, M. Furió, and P. A. Madern. 2014. The range and extent of the Vallesian Crisis (Late Miocene): new prospects based on the micromammal record from the Vallès-Penedès basin (Catalonia, Spain). J. Iber. Geol. 40.

Christiansen, P. 2008. Evolution of Skull and Mandible Shape in Cats (Carnivora: Felidae). PLoS One 3:e2807.

Cuff, A. R., M. Randau, J. Head, J. R. Hutchinson, S. E. Pierce, and A. Goswami. 2015. Big cat, small cat: reconstructing body size evolution in living and extinct Felidae. J. Evol. Biol. 28:1516–1525.

Day, E. H., X. Hua, and L. Bromham. 2016. Is specialization an evolutionary dead end? Testing for differences in speciation, extinction and trait transition rates across diverse phylogenies of specialists and generalists. J. Evol. Biol. 29:1257–1267.

de Bonis, L., S. Peigné, F. Guy, A. Likius, H. T. Makaye, P. Vignaud, and M. Brunet. 2009. A new mellivorine (Carnivora, Mustelidae) from the Late Miocene of Toros Menalla, Chad. Neues Jahrb. für Geol. und Paläontologie - Abhandlungen 252:33– 54.

DeSantis, L. R. G., B. W. Schubert, J. R. Scott, and P. S. Ungar. 2012. Implications of Diet for the Extinction of Saber-Toothed Cats and American Lions. PLoS One 7:e52453.

Eronen, J. T., M. Fortelius, A. Micheels, F. T. Portmann, K. Puolamaki, and C. M. Janis. 2012. Neogene aridification of the Northern Hemisphere. Geology 40:823– 826.

Faurby, S., M. Morlo, L. Werdelin, and K. Lyons. 2021. CarniFOSS: A database of the body mass of fossil carnivores. Glob. Ecol. Biogeogr. 30:1958–1964.

Figueirido, B., A. Martín-Serra, Z. J. Tseng, and C. M. Janis. 2015. Habitat changes and changing predatory habits in North American fossil canids. Nat. Commun. 6:7976.

Kumar, S., M. Suleski, J. M. Craig, A. E. Kasprowicz, M. Sanderford, M. Li, G. Stecher, and S. B. Hedges. 2022. TimeTree 5: An Expanded Resource for Species Divergence Times. Mol. Biol. Evol. 39.

Lisiecki, L. E., and M. E. Raymo. 2005. A Pliocene-Pleistocene stack of 57 globally distributed benthic δ 18 O records. Paleoceanography 20:n/a-n/a.

Meachen-Samuels, J., and B. Van Valkenburgh. 2009. Craniodental indicators of prey size preference in the Felidae. Biol. J. Linn. Soc. 96:784–799.

Mondanaro, A., S. Castiglione, M. Melchionna, M. Di Febbraro, G. Vitagliano, C. Serio, V. A. Vero, F. Carotenuto, and P. Raia. 2017. Living with the elephant in the room: Top-down control in Eurasian large mammal diversity over the last 22 million years. Palaeogeogr. Palaeoclimatol. Palaeoecol. 485:956–962.

Paradis, E., and K. Schliep. 2019. ape 5.0: an environment for modern phylogenetics and evolutionary analyses in R. Bioinformatics 35:526–528.

PBDB. 2023. The Paleobiology Database.

Piras, P., L. Maiorino, L. Teresi, C. Meloro, F. Lucci, T. Kotsakis, and P. Raia. 2013. Bite of the Cats: Relationships between Functional Integration and Mechanical Performance as Revealed by Mandible Geometry. Syst. Biol. 62:878–900.

Piras, P., D. Silvestro, F. Carotenuto, S. Castiglione, A. Kotsakis, L. Maiorino, M. Melchionna, A. Mondanaro, G. Sansalone, C. Serio, V. A. Vero, and P. Raia. 2018. Evolution of the sabertooth mandible: A deadly ecomorphological specialization. Palaeogeogr. Palaeoclimatol. Palaeoecol. 496:166–174.

Pires, M. M., D. Silvestro, and T. B. Quental. 2015. Continental faunal exchange and the asymmetrical radiation of carnivores. Proc. R. Soc. B Biol. Sci. 282.

Pires, M. M., D. Silvestro, and T. B. Quental. 2017. Interactions within and between clades shaped the diversification of terrestrial carnivores. Evolution (N. Y). 71:1855–1864.

Porto, L. M. V, R. S. Etienne, and R. Maestri. 2023. Evolutionary radiation in canids following continental colonizations. Evolution (N. Y). 77:971–979.

Pyron, R. A. 2015. Post-molecular systematics and the future of phylogenetics. Trends Ecol. Evol. 30:384–389.

Quental, T. B., and C. R. Marshall. 2010. Diversity dynamics: molecular phylogenies need the fossil record. Trends Ecol. Evol. 25:434–441.

R Development Core Team. 2023. R: A Language and Environment for Statistical Computing. R Foundation for Statistical Computing. Vienna, Austria.

Rabosky, D. L. 2010. Primary Controls on Species Richness in Higher Taxa. Syst. Biol. 59:634–645.

Randau, M., C. Carbone, and S. T. Turvey. 2013. Canine Evolution in Sabretoothed Carnivores: Natural Selection or Sexual Selection? PLoS One 8:e72868.

Revell, L. J. 2012. phytools: an R package for phylogenetic comparative biology (and other things). Methods Ecol. Evol. 3:217–223.

Sanderson, M. J. 2002. Estimating Absolute Rates of Molecular Evolution and Divergence Times: A Penalized Likelihood Approach. Mol. Biol. Evol. 19:101– 109.

Silvestro, D., A. Antonelli, N. Salamin, and T. B. Quental. 2015. The role of clade competition in the diversification of North American canids. Proc. Natl. Acad. Sci. 112:8684–8689.

Silvestro, D., N. Salamin, and J. Schnitzler. 2014. PyRate: a new program to estimate speciation and extinction rates from incomplete fossil data. Methods Ecol. Evol. 5:1126–1131.

Slater, G. J. 2015. Iterative adaptive radiations of fossil canids show no evidence for diversity-dependent trait evolution. Proc. Natl. Acad. Sci. 112:4897–4902.

Slater, G. J., and A. R. Friscia. 2019. Hierarchy in adaptive radiation: A case study using the Carnivora (Mammalia). Evolution (N. Y). 73:524–539.

Slater, G. J., and B. Van Valkenburgh. 2009. Allometry and performance: the evolution of skull form and function in felids. J. Evol. Biol. 22:2278–2287.

Slater, G. J., and B. Van Valkenburgh. 2008. Long in the tooth: evolution of sabertooth cat cranial shape. Paleobiology 34:403–419.

Springer, M. S., N. M. Foley, P. L. Brady, J. Gatesy, and W. J. Murphy. 2019. Evolutionary Models for the Diversification of Placental Mammals Across the KPg Boundary. Front. Genet. 10.

Sunquist, M., and F. Sunquist. 2002. Wild Cats of the World. University of Chicago Press.

Van Valkenburgh, B. 1989. Carnivore Dental Adaptations and Diet: A Study of Trophic Diversity within Guilds. Pp. 410–436 in Carnivore Behavior, Ecology, and Evolution. Springer US, Boston, MA.

Van Valkenburgh, B. 2007. Deja vu: the evolution of feeding morphologies in the Carnivora. Integr. Comp. Biol. 47:147–163.

Van Valkenburgh, B., M. W. Hayward, W. J. Ripple, C. Meloro, and V. L. Roth. 2015. The impact of large terrestrial carnivores on Pleistocene ecosystems. Proc. Natl. Acad. Sci. 113:862–867.

Wang, X., and R. Tedford. 2008. Dogs: Their Fossil Relatives and Evolutionary History. 1st ed. Columbia University Press.

Werdelin, L., and S. Peigné. 2010. Cenozoic Mammals of Africa. University of California Press, Berkeley, Berkeley, CA.

Werdelin, L., N. Yamaguchi, Warren E Johnson, and S.J. O’Brien. 2010. Phylogeny and evolution of cats (Felidae). P. in Biology and Conservation of Wild Felids.

Wiens, J. J., C. A. Kuczynski, T. Townsend, T. W. Reeder, D. G. Mulcahy, and J. W. Sites. 2010. Combining Phylogenomics and Fossils in Higher-Level Squamate Reptile Phylogeny: Molecular Data Change the Placement of Fossil Taxa. Syst. Biol. 59:674–688.

Wilson, E., and A. Mittermeier. 2009. Handbook of Mammals of the World: Carnivores. Lynx Edicions.

Zachos, J. C., G. R. Dickens, and R. E. Zeebe. 2008. An early Cenozoic perspective on greenhouse warming and carbon-cycle dynamics. Nature 451:279–283.

